# Neutral auditory words immediately followed by painful electric shock show reduced next-day recollection

**DOI:** 10.1101/2020.02.24.963165

**Authors:** Caroline M. Norton, James W. Ibinson, Samantha J. Pcola, Vencislav Popov, Joshua J. Tremel, Lynne M. Reder, Julie A. Fiez, Keith M. Vogt

## Abstract

In this study, we investigated the effect of experimentally delivered acute pain on explicit and implicit memory. Twenty-five subjects participated in experimental sessions on consecutive days. The first session involved a categorization task intended to induce incidental memory encoding. There were two conditions, presented in randomized order, in which subjects listened to a series of words, which was repeated three times. In one condition, one-third of the word items were immediately followed by a painful electrical shock and these pain-paired items were presented unpredictably. In the other condition, all word items were not associated with pain. Response times over these repeated presentations were assessed for differences. Explicit memory was tested the following day, employing a Remember-Know assessment of word recognition, with no shocks employed. Recollection was significantly reduced for pain-paired words, as the proportion of correct Remember responses (out of total correct responses) was significantly lower. There were no significant reductions in memory for non-pain items that followed painful stimulation after a period of several seconds. Consistent with the experience of pain consuming working memory resources, we theorize that painful shocks interrupt memory encoding for the immediately preceding experimental items, due to a shift in attention away from the word item.

## Introduction

This study describes the effects on long-term memory for neutral experimental items, depending on their pairing with and proximity to painful stimulations. Previous work on recognition memory for experimental items directly paired with painful stimuli have yielded divergent results. One study employing visual items, thermal pain, and immediate memory testing showed a reduction in recognition for pain-paired images (Forkmann, Schmidt, Schultz, Sommer, & Bingel, 2016). Another study using painful electric shock and images of scenes showed no effect on immediate recognition memory testing (Schwarze, Bingel, & Sommer, 2012). However, in a separate cohort (who notably performed the experiment in an MRI), next-day memory testing showed enhanced familiarity for scenes paired with painful shock (Schwarze et al., 2012). This controversy could possibly be explained by a number of experimental differences, and indicates a need for an expanded range of paradigms to better define which effects are consistent. More generally, the effect of acute somatic pain on encoding into long-term declarative memory is a complicated phenomenon with conflicting results that warrant additional investigatory attention (Vogt et al., 2019).

Our previous study (Vogt et al., 2019) employed two main experimental conditions, the first in which there was no painful stimulation following a word (No Pain Alone words). In the other condition, half of the words were immediately followed by a painful shock (Pain Mixed words) alternated with words not followed by a shock (No Pain Mixed words). Both word types in the Pain Condition showed less recollection than words in the No Pain Condition. Interestingly, a greater decrease in recollection was shown for the words that followed painful stimulation after several seconds, rather than words that were immediately followed by pain. This result demonstrated that the effect of pain may generalize to affect memory for non-pain items presented in the same experimental context. We postulated that this effect was mediated by a consumption of working memory resources from previous shocks paired with the preceding words. However, our previous study had an alternating design, and thus predictable patternof shocks, limiting the inferences that could be drawn. While Pain Mixed words were immediately followed by a shock, each No Pain Mixed word was preceded by a shock (albeit at a spacing of several seconds). Since the effect of pain on memory could extend in both temporal directions, it was unclear to what extent predictability versus proximity affected the results.

The study presented here employed a modified experimental design including 30 pain-paired words which were randomly presented among 60 non-pain items. Memory for both of these word types was compared to words presented in a totally pain-free control condition. This design effectively made shocks unpredictable and less frequent. As a result, while the pain-paired words were followed by pain, as in our previous design, the no-pain paired words were no longer systematically preceded by a shock. We also conducted the long-term recognition memory testing the next day (rather than immediately afterwards). This longer follow-up interval, coupled with the increased number of experimental items, was anticipated to decrease the impact of any ceiling effects on memory performance. We hypothesized that the interruptive effect on No Pain Mixed items would be reduced, and that this would result in a significant decrease in explicit memory (particularly recollection) for Pain Mixed word items, compared to both types of non-pain words.

As a secondary outcome, we sought to determine if pain-pairing would elicit priming effects observable in reaction time (RT) measurements during encoding, as these could indicate implicit memory for the pain pairing, even in the absence of long-term recognition memory results. The design included three repetitions of each word list, so changes in RT over the course of the experiment could be assessed. Similar to our previous results, which showed overall slower response times for words in an experimental context containing painful stimulation (Vogt et al., 2019), we predicted Pain Mixed and No Pain Mixed words would have longer response times than No Pain Alone words. An additional secondary outcome was the effect that pain would have on memory for both the current and subsequent words. We expected to see a decrease in recollection for words that more often immediately followed a pain stimulation, with less of an effect for non-pain words that followed pain words less frequently.

## Methods

### Subjects

#### Power Analysis

We used our previously reported data (Vogt et al., 2019) to determine an adequate sample size for the present study. Specifically, the difference in d′ detected in the previous study between the No Pain Alone and No Pain Mixed word types was 0.33, with the variance of that difference being .30. Using these estimates and SamplePower 3.0.1 (IBM, New York, NY), a sample size of 25 subjects was estimated to have 82% power to detect a difference this large for this primary outcome. This calculation assumes paired data (from the same subjects) and alpha = .05 (2-tailed). Applicability of this power analysis to the present study also assumes that despite experimental differences in study design, the present study will have a similar mean and variance as compared to our past study.

#### Demographics

Data were acquired from healthy volunteer subjects between the ages of 18 and 30 who were recruited from the university community. They received $10 per hour as compensation. Eligibility was determined by self-report of exclusionary criteria. All subjects acknowledged being free from significant memory impairment, hearing loss, sleep apnea, chronic pain, other chronic medical problems, neurologic and psychiatric diseases, as well as the use of antidepressants, antipsychotics, antihistamines, antianxiety medication, stimulants, sleep aids, and pain medication. Data presented are from a cohort of 25 subjects (15 female) with age in years 22.0 ± 3.2 (mean ± standard deviation). No subjects were lost to follow-up and none had entirely unusable data. The study was approved by the University of Pittsburgh Institutional Review Board (PRO16110197) and conformed to all relevant standards for the ethical and responsible conduct of research.

### Procedures

#### Encoding Task Procedure and Design

After informed consent, subjects were given task instructions and an electric nerve stimulator was titrated to a subjective pain rating of 7/10 on a numerical scale from 0-10, anchored with 0 being none and 10 being the worst imaginable. The current necessary to achieve this target pain rating was determined, and subjects underwent a short practice session during which they briefly experienced the pain stimulus two more times. Following the practice, subjects again rated their pain experience and the nerve stimulator was re-adjusted, if desired by subjects, to the target a 7/10 rating, followed by repeating the practice session. After this point, the intensity of the nerve stimulator was not manipulated in the remainder of the experiment. A second pain rating was obtained after the pain condition, but pain ratings for individual items were not obtained during the experiment.

The design of the experiment is depicted graphically in Figure 1. The two main portions of the study were the encoding portion and the memory testing portion. During the encoding portion, subjects listened to a series of words and made decisions about them. The words used for this task were the same auditory recordings used in our previous study (Vogt et al., 2019), consisting of commonly-used non-proper nouns. There were two conditions within the encoding portion of the study: the Pain condition during which one third of the words were immediately followed by a 1-second electric shock, and the No Pain condition, during which no pain was experienced for any word. Subjects were informed by onscreen labels which condition they were about to experience, and thus warned in advance whether the coming experimental period would include painful stimulation. Each of the two conditions consisted of a list of 90 words repeated three times, in random order. The pain-paired words and non-pain words in the Pain condition were distributed with the constraint that no more than two pain-paired words occurred consecutively, and no more than five non-pain words occurred consecutively. Individual recorded words were presented through headphones, and subjects made decisions about them while seated at and interacting with a laptop computer. Subjects were asked to judge each word on a particular dimension, the decisions used were: (1) moves or not, (2) living or not, (3) natural or not, (4) would fit in a shoebox or not, (5) place or event or not, and (6) typically used or not. Three decisions were assigned randomly to each of the two encoding conditions. Subjects were able to respond any time after the start of the word being played, as well as during the subsequent shock, if one occurred. The maximum response time (RT) window was six seconds, after which the next word would be presented automatically. The assignment of a set of decisions to a condition (Pain vs. No Pain), the order of the decision list within a condition, and the order of the words within a decision list were randomized for each subject, with the constraint that half of the subjects received the Pain Condition first.

**Figure 1.**
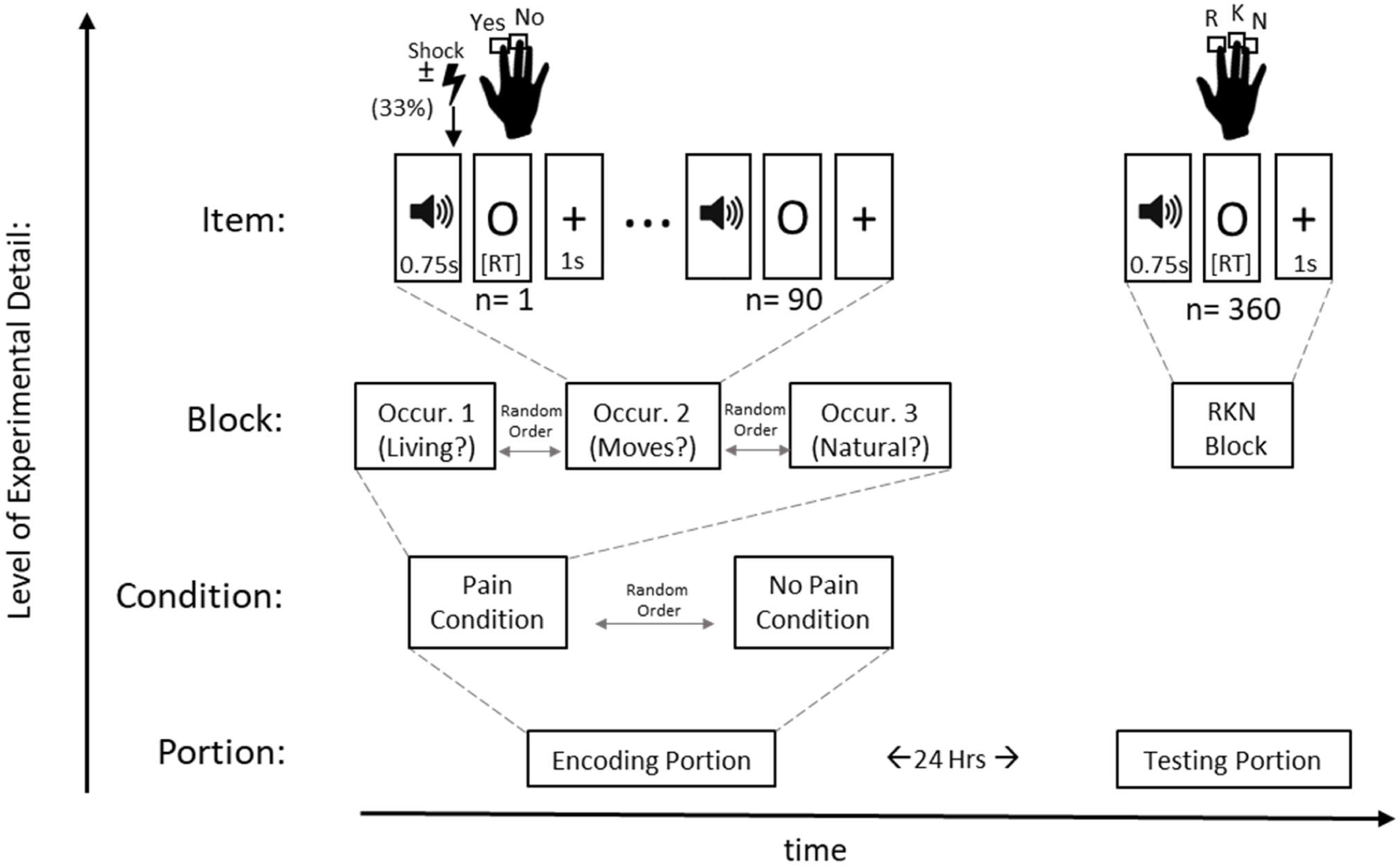
Graphical depiction of experimental design in nested layers of detail, labelled on the left. Time is represented along the x-axis. Increasing level of experimental detail is shown going from bottom to top, with dashed lines indicating expanded detail of items within a larger hierarchical part of the experiment. The two main portions of the experiment, Encoding and Testing, are shown as blocks at the bottom, these always occurred consecutively. The Encoding portion has two conditions: Pain and No Pain, and the order of these was randomized. Each Encoding condition consisted of three repetitions of a word list, examples of which are shown expanded above the Pain condition. Both Pain and No Pain conditions contained 3 occurrences of the same 90-word list. As shown in the Item level of detail, words were delivered auditorily, over 0.75 s. In the Pain Condition (only) a painful electric shock immediately followed one third of the words (30 out of 90). No shocks occurred in the No Pain condition or in the Testing portion of the experiment. Abbreviations: RT= reaction time, RKN= Remember, Know, New, Alt= alternating.

#### Memory Testing

Explicit memory testing occurred the day following the encoding portion. Recognition testing was employed, using the well-known Remember-Know-New (RKN) scheme; for a recent review, see (Migo, Mayes, & Montaldi, 2012). Subjects were given a printed sheet describing the Remember, Know, and New responses for the RKN procedure and were asked to read it. These instructions parallel those established by previous investigations (Rajaram, 1993), and are identical to the instructions available as supplementary materials to our previous publication (Vogt et al., 2019). To ensure comprehension of the task, in particular the distinction between Remember and Know responses, subjects were asked to explain the RKN procedure back to the investigator, who corrected any misunderstandings. A standardized recording summarizing these instructions was also played to the subject before beginning the RKN test. All the words heard during the previous day were played intermixed with an equal number of foils (360 items total). Word order was randomized, and each word drawn from the bank had equal probability of being assigned as a foil, versus used in the experiment. No shocks were delivered during the RKN testing session, which was announced to subjects in advance.

### Equipment

Electronic recording files previously made were used for the word stimuli (Vogt et al., 2019). Pain stimuli were generated using the same electric nerve stimulator as our previous study, with identical placement of the electrodes (Vogt et al., 2019). Data were digitized using a BIOPAC MP160 (BIOPAC Systems, Goleta, CA) data acquisition unit and Acqknowledge version 5.0 (BIOPAC Systems, Goleta, CA), running on a Windows 10 laptop PC. All parts of the experiment were implemented with E-Prime version 2.0 (Psychology Software Tools, Sharpsburg, PA), and E-Prime captured response time data. Shock delivery synchronization to follow experimental word items was accomplished using custom hardware, as in our prior study (Vogt et al., 2019) that allowed E-Prime to control the nerve stimulator through a USB connection.

### Data Analyses

For the analyses, items from the encoding portion of the experiment were categorized into three groups. Pain Mixed words were the 30 words paired with a painful shock in the Pain Condition. No Pain Mixed words were the 60 words in the Pain condition that were not paired with pain. No Pain Alone words were the 90 words experienced in the No Pain condition (which was completely without noxious stimulation).

All statistical analysis was carried out in SPSS Statistics 23 (IBM, New York, NY) with P = .05 as the threshold for significance. Bonferroni correction for multiple comparisons was used for comparing main effects. For efficiency in data manipulation, outlier removal was carried out in RStudio (version 1.0.153, https://www.rstudio.com/) using R version 3.2.5. As in our previous study (Vogt et al., 2019), the Median Absolute Deviation (MAD) was calculated, and response times more than 3.5 MAD from the median were defined as outliers and removed. We also removed incorrect RKN responses from RT analysis. After removal of these extraneous data, the common logarithm (log10) was used to transform the RT data to a normal distribution. All RT analysis was performed on the transformed data, shown in Supplementary Figures 1 and 2. For the encoding RT data, a linear mixed model (LMM) analysis was used with compound symmetry covariance structure. Occurrence and word type were selected as repeated fixed factors, and condition order was selected as an additional fixed factor. Transformed RKN RT data were also analyzed using a 2-way 3×2 Analysis of Variance (ANOVA) with word type and condition order as separate factors. Though all RT analyses were performed on transformed data, the RT figures are shown in milliseconds to aid in meaningfully visualizing the data.

Explicit memory was evaluated using signal detection theory to estimate memory sensitivity. This analysis should eliminate individual subjects’ bias toward identifying a word as remembered (vs. novel) by considering both hit and false alarm responses. Hits were counted as previously heard words correctly identified as such as either Remember or Know. False alarms were foils (words not previously heard) that were identified by subjects (as appearing in the previous part of the experiment) with either a Remember or Know response. To calculate d′, we used the formula z(hits) – z(false alarms), with z() referring to the cumulative Gaussian distribution function. To separate memory strength based on Remember versus Know responses, Remember only d′ was calculated using only correct Remember response hits and only Remember response false alarms. Some subjects achieved a perfect hit rate or zero false alarm rate, neither of which is calculable in the formula above for d′. In these cases, the estimated hit and false alarm rates were adjusted based on the assumption that there would have been one miss or one false alarm response if there twice as many items had been tested.

Familiarity was quantified, accounting for the fraction of words already identified as recollected. We employed the same adjusted familiarity score as our previous study (Vogt et al., 2019), using the previously-described formula (Yonelinas, Aly, Wang, & Koen, 2010): (Know Hit Rate – Know False Alarm Rate)/(1 – Recollection), where Recollection = Remember Hit Rate – Remember False Alarm Rate. To dissect the effect of word type within each response type category, 2-way 3×2 ANOVAs were run for Remember and Know, Remember Only, and Know Only (with Know Only represented by adjusted familiarity score) separately with word type and condition order as the independent factors.

In addition to calculating d′, memory strength was also assessed by calculating the proportion of Remember responses out of total hits. The number of correct Remember responses for each word type was tabulated and divided by the total number of hits (Remember and Know responses combined) for that word type, for each subject. These values were then compared across all subjects using a 2-way 3×2 ANOVA with word type and condition order as the independent factors.

Finally, we undertook an analysis to compare the effects of pain stimulations on memory for the current word and subsequent words. This bidirectional analysis is illustrated in Figure 2. Because of the randomized repetition structure of the experiment, non-pain words did not consistently appear in fixed position relative to pain-paired words. Post-hoc categorization of No Pain Mixed words with their relative frequency of following a pain-paired word was necessary. Memory for No Pain words was assessed, based on the percentage of times they followed a pain stimulus. Because relative word order varied with each of the three word list repetitions within the Pain Condition, data was binned based on the proportion of times that the non-pain word item followed a pain-paired word (and thus followed several seconds after pain stimulation). Because the pain-order analysis relates only to effects during the Encoding Portion of the experiment, and the foils are shared across words from each pain-order, hit rate was used as the independent variable (rather than d′). Thus, each word fell into one of four categories: 0% (never followed a Pain Mixed word), 33% (followed a Pain Mixed word for one of the three trials), 67% (followed a Pain Mixed word for two of the three trials) and 100% (followed a Pain Mixed word for all three trials). This categorization was performed for both Pain Mixed and No Pain Mixed words. To assess for differences between the four percentage categories within individual response types (Remember only vs. Remember + Know) as well as within the Pain Mixed and No Pain Mixed word types, one-way ANOVA analyses were conducted for hit rates within each sub-category (i.e., Remember responses for No Pain Mixed words, Remember and Know combined responses for Pain Mixed word, etc.). To see if following a Pain Mixed word on any of the occurrences had an effect, paired T-tests were used to compare words that never followed a pain stimulation (labelled “0%”) to the average across the three nonzero proportions of following a pain stimulation (collectively labelled “>0%”).

**Figure 2.**
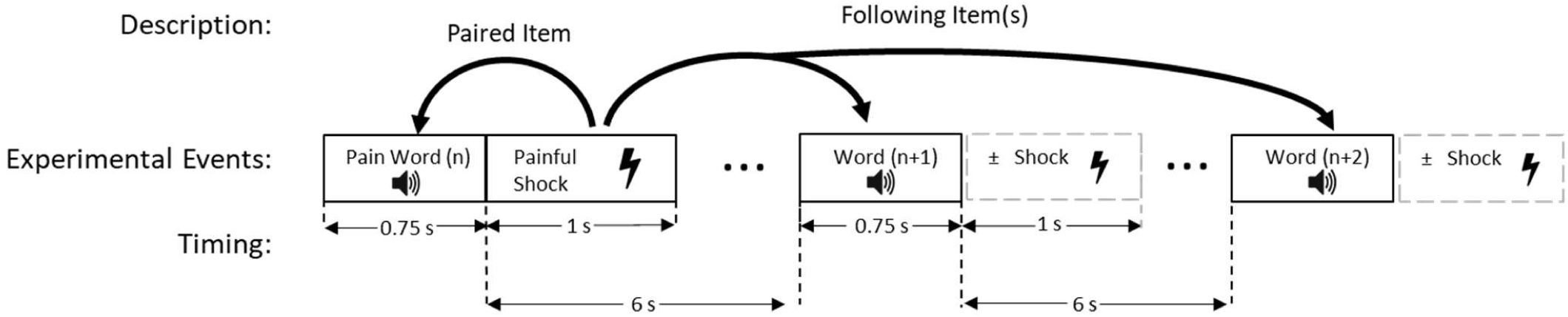
Graphical depiction of temporal bidirectionality of the effect painful shocks may have on memory. Paired items refer to the Pain word that the shock immediately follows. Following items, labelled n+1 and n+2, may be pain-paired or non-pain words that may have their own immediately-associated shock events. Abbreviation: s= seconds

A second pain-order analysis examined the memory results based only on the relative order in the first word list (and thus the first occurrence of each individual word). This analysis is based on the assumption that the pain context of the first presentation may have a much stronger effect on memory encoding than any effects during the second and third presentations. Pain-paired words are labelled item n, and the subsequent item labelled n+1. To account for potential differences based on word order within each response type and word type, paired T-tests were conducted such that the n and n+1 Hit Rates were compared for Pain words within each Response Type, and n + 1 and n + 2 or more words were compared for No Pain Mixed words within each response type.

## R esults

### Pain Intensities

Nerve stimulator intensity, as selected by the subjects to be a 7 out of 10 pain rating at the start of the experiment, was 11.4 ± 4.7 mA. Subjective pain ratings obtained just after the practice session were 6.1 ± 1.0. Pain ratings following the Pain condition were 6.0 ± 1.0.

### Response Time

Histograms showing RT values before and after transformation are shown in Supplementary Figures 3 and 4. Response time data are presented with outliers removed. For the RT data, outliers were detected in 4.1% of responses during the encoding portion and 4.6% of RKN responses.

Response times for the three word types across the three repetitions in the encoding portion are shown in Figure 3. Only one response time for one subject was excluded for being longer than the pre-determined response window limit of 6 seconds. The RTs (in ms) collapsed across all three word types and both condition orders during occurrence 1 (mean = 1368, 95% CI = 1285 - 1450) were significantly greater than occurrence 2 (P < .001, mean = 1284, 95% CI = 1201 - 1366) as well as occurrence 3 (P < .001, mean = 1260, 95% CI = 1178 - 1343). Response times for occurrence 3 were not significantly different from those for occurrence 2 (P = .57). Response times for the No Pain Alone words (mean = 1304, 95% CI = 1221 - 1386) were not significantly different from No Pain Mixed words (mean = 1305, 95% CI = 1223 - 1387, P = 1.00) or Pain Mixed words (mean = 1303, 95% CI = 1221 - 1385, P = 1.00) when averaged across all occurrences and both condition orders. Similarly, response times for No Pain Mixed words were also not significantly different from those for Pain Mixed words (P = 1.00). Though not shown graphically, response times for subjects who experienced the Pain condition first (mean = 1323, 95% CI = 1208 - 1438) were not significantly different from those who experienced the No Pain condition first (mean = 1285, 95% CI = 1174 - 1395, P = .55) when collapsed across all occurrences and word types. There were also no significant interaction effects between word type and occurrence.

**Figure 3.**
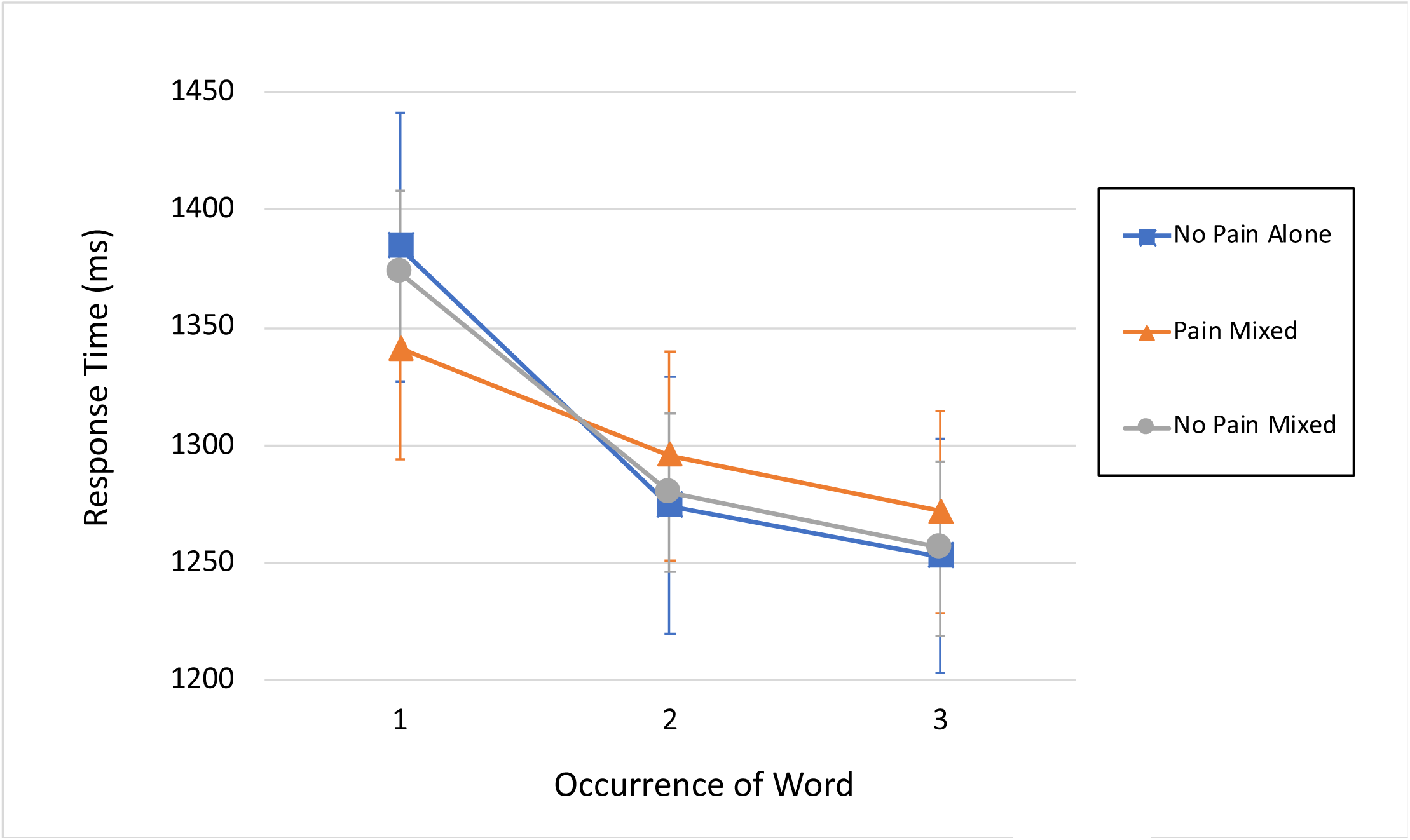
Response times over the three encoding repetitions for each word-type. Error bars represent standard error.

The RKN recognition memory task RT data are shown in Figure 4, limited to correct responses only. Correct responses represented 91% of total responses for Remember, 68% of total responses for Know, and 83% of total responses for New. As expected, Know response times (mean = 1775, 95% CI = 1671 - 1879 were significantly longer than Remember response times (mean = 1531, 95% CI = 1428 - 1635), when collapsed across all three word types (P < .001). When comparing within Remember and Know responses types, there were no significant differences among the three different word types. Though the data are not shown divided this way, there were no significant RT differences between the two conditions orders. Response times for words correctly identified as New were significantly shorter than Know responses (P < 0.001).

**Figure 4.**
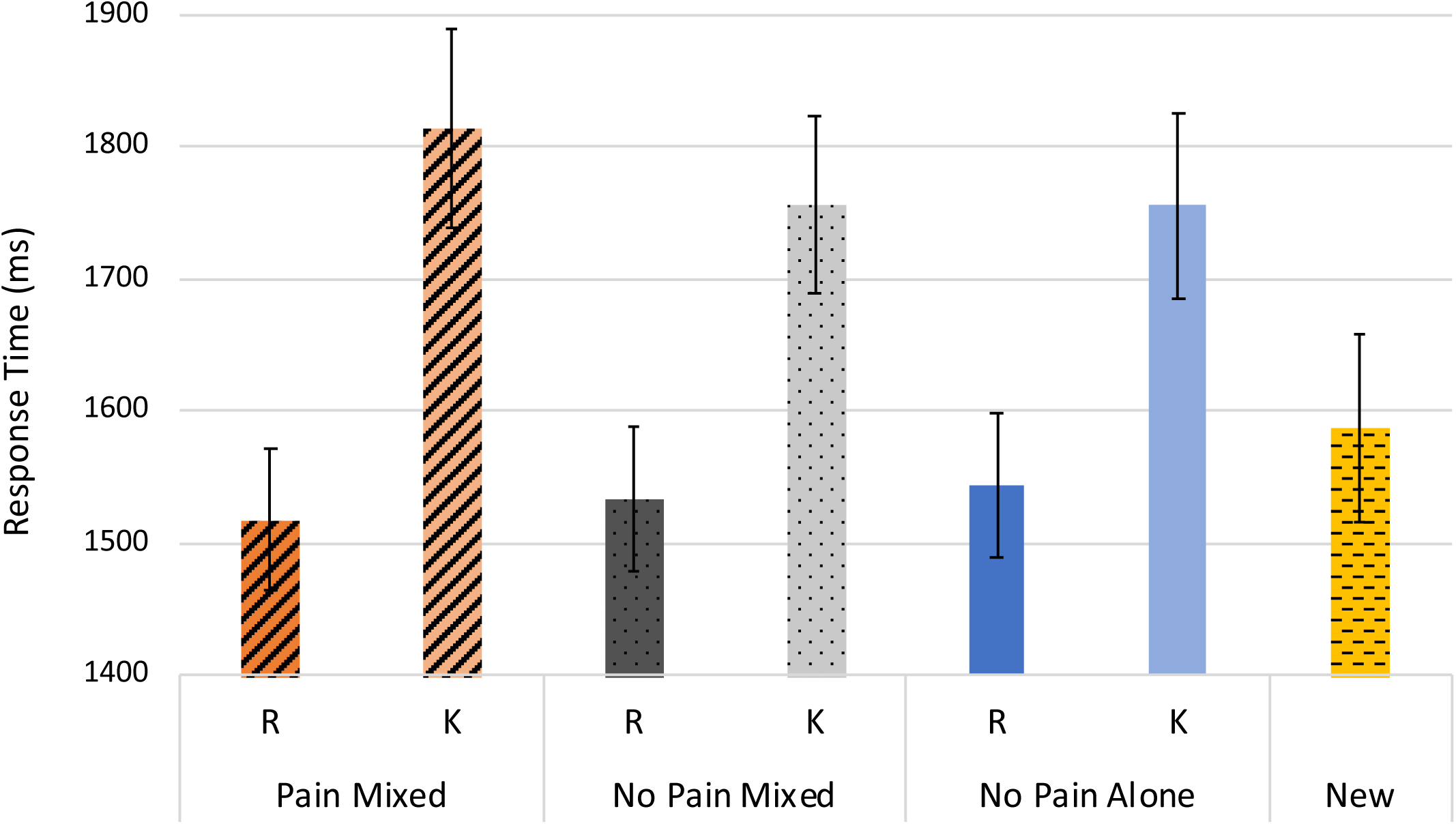
Response times for Remember (R), Know (K), and New (N) responses during the RKN testing portion, including only correct responses. Error bars represent standard error. Significant differences reported in Results section.

### Memory Testing

The d′ values calculated for combined Remember and Know (R+K) responses, and Remember responses alone are shown in Figure 5A. For transparency, the hit rate and the false alarm rates for each subject, from which these are calculated, are shown in Supplementary Table 1. Examining combined R+K responses, d′ for No Pain Alone words (mean = 2.15, 95% CI = 1.91 - 2.38) were not significantly different (P = .47) from Pain Mixed words (mean = 1.91, 95% CI = 1.68 - 2.15) nor from No Pain Mixed words (mean = 2.11, 95% CI = 1.89 - 2.35, P = 1.00). Pain Mixed words also did not have significantly different d′ (P = .71) than No Pain Mixed words for combined R+K responses. Thus, there was no main effect for word type. There was a significant main effect for condition order, where subjects who experienced the Pain condition first had significantly higher (P = .008) combined R+K d′ scores (mean = 2.24, 95% CI = 2.05 - 2.44) than those who experienced the No Pain condition first (mean = 1.87, 95% CI = 1.68 - 2.06). There was no significant interaction between word type and condition order.

**Figure 5.**
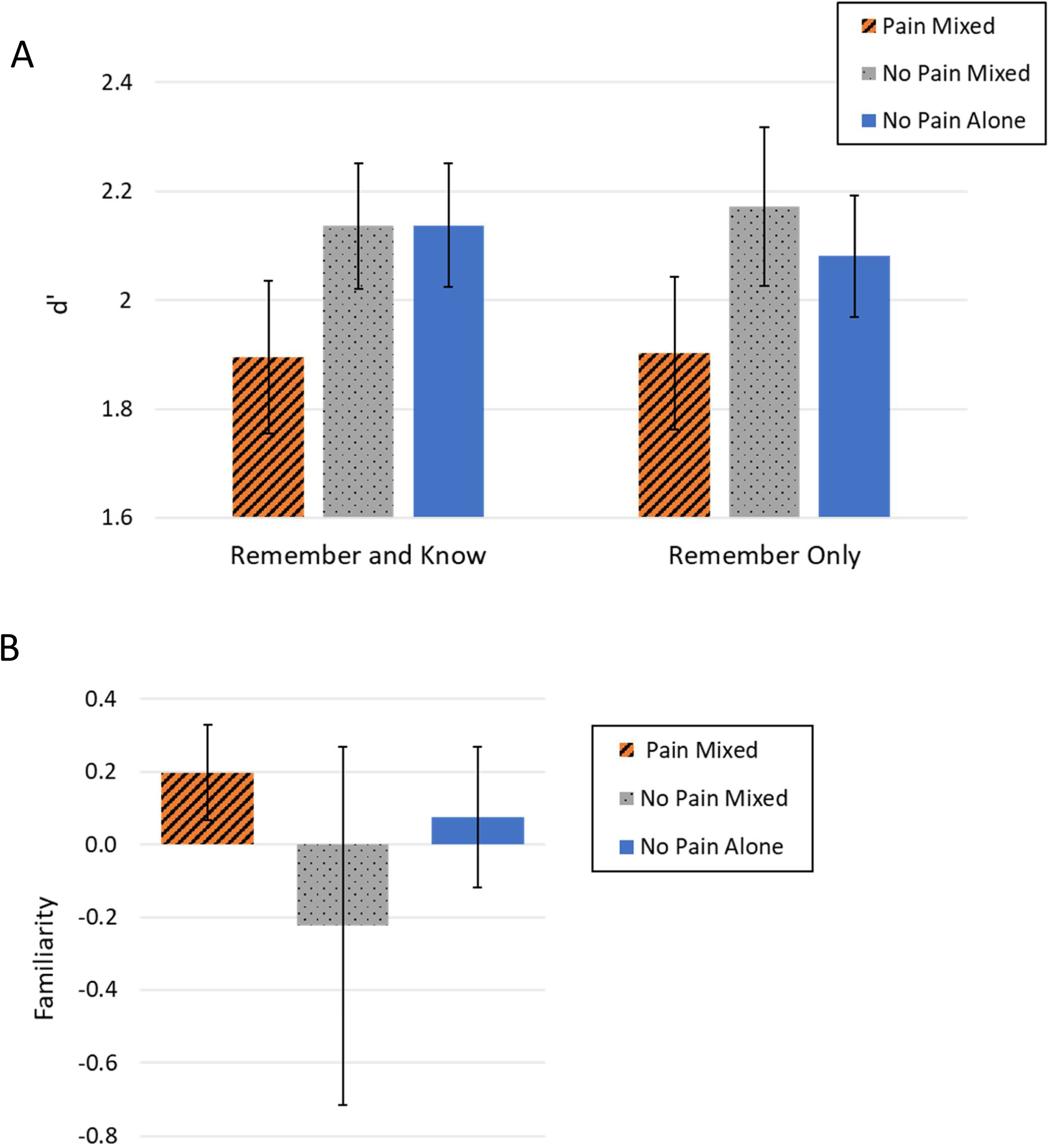
Recognition memory performance across word-type. Panel A compares average d′ values for overall recognition (combined Remember and Know responses) and specifically for recollection and familiarity (Remember and Know responses separately). Panel B compares a composite measure of familiarity (see text for details). Error bars display standard error. No differences were statistically significant.

For Remember Only responses, the d′ scores for No Pain Alone words (mean = 2.07, 95% CI = 1.81 - 2.34) were not significantly different (P = 1.00) from Pain Mixed words (mean = 1.90, 95% CI = 1.64 - 2.17) or No Pain Mixed words (mean = 2.17, 95% CI = 1.91 - 2.44, P = 1.00). The d′ difference between Pain Mixed words and No Pain Mixed words was also not significantly different for Remember Only responses (P = .47). Hence, there was no main effect for word type. Subjects who experienced the Pain condition first did not have significantly different d′ scores (mean = 2.01, 95% CI = 1.80 - 2.23) from those who had the No Pain condition first (mean = 2.09, 95% CI = 1.88 - 2.30), so there was also no main effect for condition order for Remember responses. The interaction between word type and condition order was also not significant (P = .46). The adjusted familiarity scores are shown in Figure 5B. For this parameter, No Pain Alone words (mean = .07, 95% CI = –.55 - .67) did not have significantly different scores (P = 1.00) than Pain Mixed words (mean = .19, 95% CI = –.42 - .80) or No Pain Mixed words (mean = –.25, 95% CI = –.86 - .36, P = 1.00). The familiarity scores for Pain Mixed and No Pain Mixed words were also not significantly different (P = .95). There was thus no significant main effect for word type. Subjects who experienced the Pain condition first (mean = –.33, 95% CI = –.84 - .18) did not have significantly different (P = .06) familiarity scores than those who experienced the No Pain condition first (mean = .34, 95% CI = –.15 - .82), so there was also no significant main effect for condition order. The interaction between word type and condition order was also not significant for familiarity scores (P = .69).

The proportion of Remember responses out of total Remember + Know (R+K) hits is shown in Figure 6. Unlike Hit Rate alone, this measure illustrates recollection compared to familiarity in one composite score. False alarm rates were shared between the three word types, so not accounting for false alarms should not impact interpretability. This proportion of Remember responses for Pain Mixed words (mean = .47, 95% CI = .38 - .55), was significantly lower (P = .007) than No Pain Mixed words (mean = .65, 95% CI = .57 - .74) and also significantly lower (P = .04) than No Pain Alone words (mean = .62, 95% CI = .53 - .70). The proportion of Remember responses was not significantly different (P = .89) when comparing subjects who experienced the Pain condition first (mean = .58, 95% CI = .51 - .65) and those who experienced the No Pain condition first (mean = .58, 95% CI = .51 - .64), so there was no significant main effect for condition order on the proportion of Remember responses out of total hits. There was also no significant interaction between word type and condition order interaction for this parameter (P = .43).

**Figure 6.**
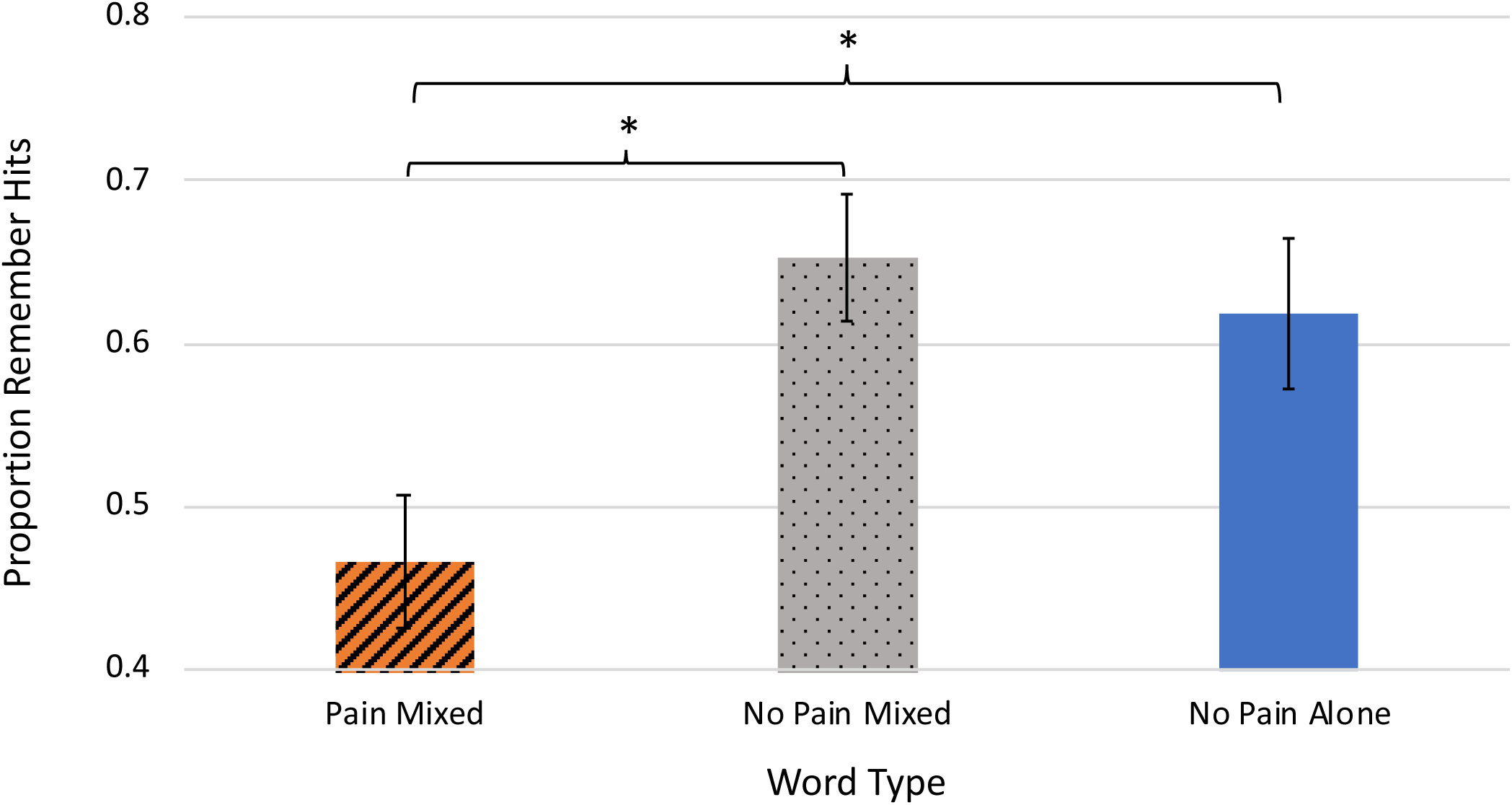
Proportion of Remember hits out of total hits for each word type. Error bars display standard error. Significant differences are indicated with an asterisk (*).

Figure 7 shows graphically the memory performance results in the pain-order analysis for combined R+K responses. Data is binned by percentage of times that a given word item followed a pain-paired item (and thus followed a pain stimulation) across the three repetition blocks, with Panel A showing all three pain-order possibilities and Panel B showing a binary split between words that never followed a pain stimulation, versus those that did at least once. Figure 8 displays data parallel to that described for Figure 7, except examining Remember responses only. One-way ANOVA analyses revealed no significant Hit Rate differences between percentage categories within any of the word type in either RKN response type sub-category. Thus, there were no significant differences between any data shown in Figure 7 or 8.

**Figure 7.**
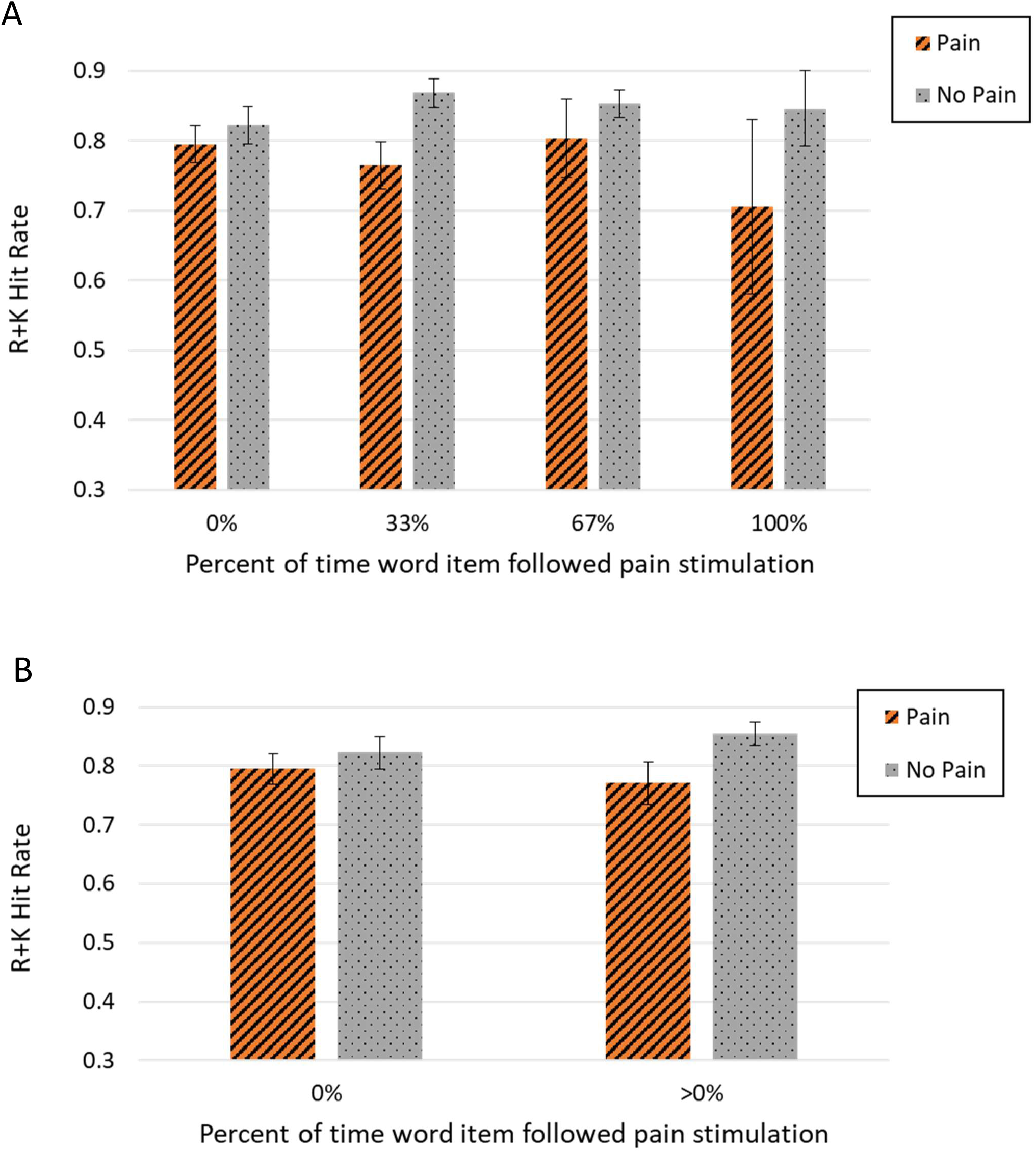
Hit rates for Remember and Know combined, separated by percent of the time that a word immediately followed a pain word. Panel A shows words that followed a pain word 33%, 67%, and 100% separately. Panel B combines all three of the nonzero percentages in Panel A into a single >0% group. Error bars display standard error. No differences were statistically significant.

**Figure 8.**
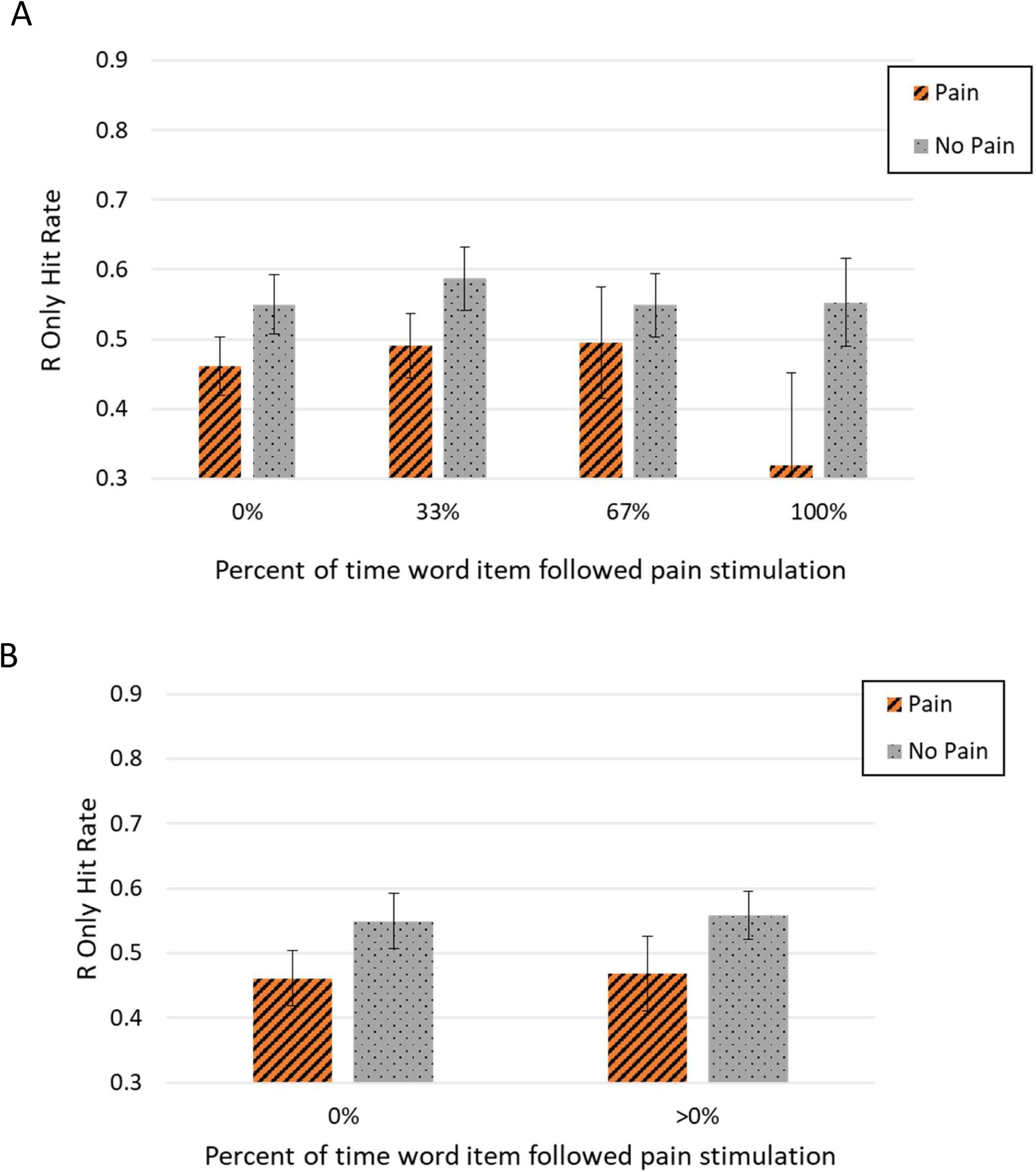
Hit rates for Remember only responses, separated by percent of the time that a word immediately followed a pain word. Panel A shows words that followed a pain word 33%, 67%, and 100% separately. Panel B combines all three of the nonzero percentages in Panel A into a single >0% group. Error bars display standard error. No differences were statistically significant.

Figure 9 shows data from the first Pain Condition trial block only, and thus the first pain-ordering experienced by subjects in the experiment. Mean hit rates for R+K (panel A) or R only (panel B) responses were not significantly different, using paired T-tests to compare data binned by this first ordering.

**Figure 9.**
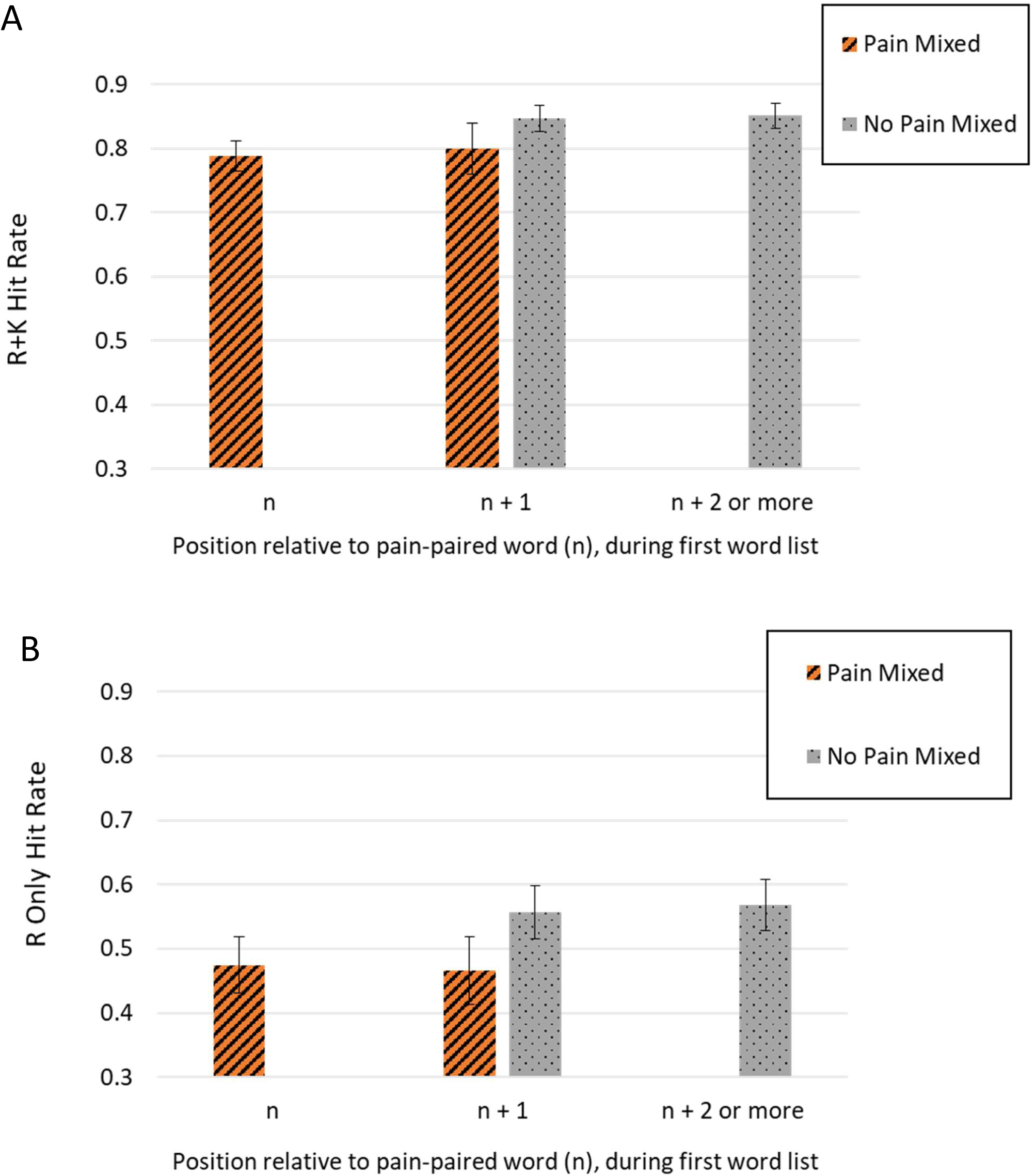
Hit rates for Remember and Know combined responses (R+K; Panel A) as well as Remember Only responses (R Only; Panel B) for words that did not immediately follow a Pain Mixed word (Not After Pain) and words that occurred immediately following a Pain Mixed word (After Pain). No differences were statistically significant.

## Discussion

The main goal of this study was to further elucidate the temporal effects of acute pain on recognition memory for auditory word items with variable proximity to painful electric stimuli. Memory was compared for three different word types; Pain Mixed words (immediately followed by a shock) and No Pain Mixed words (not paired with shock, but in the same experimental condition), as well as No Pain Alone words (heard in a completely pain free context). Our previous study using the same word items and conditions employed alternating pain-pairing, and thus a predictable pattern, and immediate RKN testing found that Remember-only d′ and the proportion of Remember (to overall) hits were significantly lower for No Pain Mixed, compared to No Pain Alone words. This result was attributed to the interruptive effect of previous pain stimulation on subsequently-occurring non-pain words. The present study had distinguishing features of unpredictable pain-pairing for one-third of the words in the Pain Condition and next-day RKN testing.

### Recognition Memory Performance

Contrary to our expectations, comparisons between measures of recollection and familiarity for Pain words, compared to both types of non-pain words, were not statistically significant. However, Pain words displayed fewer Remember responses, as a proportion of all hits (Remember + Know), compared to both types of non-pain words. This result demonstrates that, for words items immediately followed by a painful stimulus, subjects had less ability to bind the item within the experimental context. On close examination, the non-significant comparisons between d′ values in Figure 5A reflect the high variability in false positive rates. Similarly, the comparisons in Figure 5B are non-significant due to variance in the familiarity composite measure, likely driven by more variability in Know responses.

Our results agree with previous research showing that pain can impair memory performance for pain-paired experimental items when examining d′ scores for both recollection and familiarity (Forkmann et al., 2016). Importantly, subjects in that study rated their expectation of pain’s effect on memory prior to memory encoding, so the differences in memory performance could have been biased by subject’s prior expectations. We did not prospectively ask our subjects their expectations for pain’s effect on memory, so they would have been less likely to make predictions about the potential effect of pain on their memory performance during the experiment. Thus, our results for pain-induced decreases in memory should be independent of subjects’ expectation of this impairment.

To further understand the observed effect of pain on explicit memory, inference can be drawn from the literature on how modulated attention affects memory. Divided attention can impair explicit memory in a variety of settings (Amado, 2000; Kilb & Naveh-Benjamin, 2015; Liang & You, 2003; Mulligan, 1998). These studies typically involve an overt distraction task during memory encoding, such as identifying a string of aurally presented numbers while trying to encode visually-presented words (Mulligan, 1998). Subjects’ attention was not specifically manipulated or quantified in the present study. However, it is reasonable to infer that the experience of significant pain is sufficiently distracting to impact participant’s attention, leading to decreased memory for words immediately followed by (and thus paired with) a painful shock. Though performance on a working memory task was not directly examined in this study, engagement of working memory resources, necessary for successful encoding into long-term memory (Baddeley, 2010), can be reduced by other experimental events (Popov & Reder, 2018). A division of attention is consistent with a reduction in working memory resources available for encoding, specifically binding the word items to experimental context.

Our previous study (Vogt et al., 2019) showed the lowest recollection for No Pain Mixed words. Because of the alternating pattern of presentation, all of the No Pain Mixed words followed a pain stimulation, after a few second delay. To better describe these potential aftereffects from pain stimulation in the present study, we performed a secondary analysis of memory for words that followed a pain-paired word. Because the word lists were presented in different orders for each of the three repetitions, the presence of a preceding pain word was variable across the occurrences of each No Pain Mixed word. To account for this design, we calculated the percentage of time that a non-pain word followed a pain stimulation. There were four possibilities for the frequency that any word in the Pain Condition (both Pain Mixed and No Pain Mixed) followed a shock: 0%, 33%, 67%, or 100% of the time. Since there were fewer words categorized into the 67% and 100% bins, we also analyzed for effects comparing between words in two categories: 0% and the combined >0% incidence after a Pain Mixed word. With each of these comparisons, and conservative statistical analyses, no significant differences for hit rates were detected for Remember responses alone or the combined R+K responses. Finally, we analyzed memory for words grouped based on the order of presentation in the first word list only. This analysis presumes that this initial exposure may have more significantly influenced memory encoding than the combined effect of the two subsequent words presentations. Thus, we compared hit rates based on each word’s position, relative to its most proximate preceding pain-paired word. The small differences in hit rate between these groups were not statistically significant. Taken together, the reduction in recollection that painful stimulation has for an experimental item it is paired with (immediately follows) seems to outstrip any effect it has on memory for experimental items that follow the pain by a few seconds. If another experiment were performed with the constraint of consistent positioning of Pain Mixed and No Pain Mixed items relative to one another across three trials, it would be possible to directly assess the effects of painful stimulation on both words that immediately precede it and those that follow by some delay. This discrepancy illustrates the importance of considering how the timing of pain stimulation in an experiment may impact explicit memory results. We recommend that future investigations be verbose in descriptions of this timing, as this specificity is lacking in some of the previous relevant literature.

### Encoding Response Times

As expected, response times decreased for all three word types over the course of the three trials in the encoding portion of the experiment. This result is an expected effect of practice, whereby subjects are responding faster as they get more familiar with the task, and is in line with the results of our previous study (Vogt et al., 2019). However, we did not find any significant differences between word types for the encoding portion of the study, suggesting that neither pain-pairing nor pain context had an interaction effect on task response over time.

### Memory Testing Response Times

We did not observe any significant response time differences between the three different word types during the RKN memory testing task. Because longer response times can improve recognition accuracy, this consistency of response times across conditions being compared indicates that the differences observed in memory performance were not confounded by differing amounts of contemplation when subjects were determining how to respond. Moreover, longer response times for Know responses compared to Remember responses are in line with previous results for RKN testing (Rotello & Zeng, 2008). This supports that our subjects understood the RKN task, and the differences between Remember and Know responses are more reflective of differences in recollection versus familiarity, rather than a probabilistic measure of confidence.

### Limitations

We agree with the dual-process model of recognition (Reder et al., 2000), in which recollection and familiarity are viewed as occupying distinct measures of explicit memory (Diana et al. 2006), though we also recognize this theory is debated (Wais, Mickes, & Wixted, 2008). Additionally, some subjects certainly gave differing proportions of Remember versus Know judgements, which could either indicate broad variability in memory strength across subjects or differing levels of understanding the RKN task, despite the measures taken to mitigate any confusion and the constancy in RTs described above.

Our experimental framework and results can also be contexualized with other related lines of memory research that were not specifically addressed experimentally. Emotional valence was not specifically controlled for and could vary across subjects for individual word items. However, as words used in the experiment were drawn from a bank of 720 potentials, any individual memory effects from emotional valence should be evenly distributed across experimental items. As shocks occurred after each Pain Mixed word was played, any effect on emotional valence should be mostly attributed to the pain stimulation, rather than the word itself. As subjects could not predict nor directly influence the timing of painful stimulations during this experiment, motivation, punishment, or reward mechanisms were similarly not the primary cognitive mechanisms being investigated.

Though the present study did allow for assessment of temporal bidirectionality in the effects of pain on memory, word presentation order was not specifically constrained to maximize this comparison. As discussed above, word order was randomized across the three list presentations, making the relative proximity of non-pain words to painful stimuli variable between experimental blocks. Our categorization strategy for non-pain words, based on their frequency of presentation following pain words, is inherently limited in its predictive ability. This variability is exacerbated by the small number of non-pain word items for which all three presentations occurred after a word-shock pair. In fact, some subjects had no words meeting this criterion during their experiment. This limitation is why words in the 100% frequency bins in Figures 7 - 9 have great variance and may not reliably represent any ordering effects from previous pain stimulation. Though they are more numerous, words in the 33% and 66% frequency bins would inherently have a diluted influence from any pain versus non-pain sequence effects.

Finally, other experimental design choices could have also influenced the results. It is reasonable to presume that, with fewer pain-paired items and an unpredictable pattern, the interruptive effect of a shock on subsequent (non-pain) items would have less of an impact on memory encoding. The less frequent occurrence of painful stimulations could also be expected to reduce the overall averseness of the Pain Condition, but this was not specifically quantified by subject report in either of our studies. However, it is reasonable to presume that the experimental framework of the study reported here is more focused on the direct effect of pain on item memory, rather than any general effects from an overall painful experimental context.

## C onclusions

Employing an experimental framework with an unpredictable pattern of periodic pairing of auditory word items with painful electric shock, we used next-day recognition RKN testing to determine pain’s effect on memory. Recollection for words immediately followed by shock was reduced, compared to two groups of non-pain words: those heard intermixed with pain-paired words and also those heard in a completely pain-free context. Consistent with the experience of pain consuming working memory resources, we postulate that painful shocks interfere with memory encoding for immediately preceding experimental items, due to a shift in attention away from the word item. This study did not demonstrate memory reduction for non-pain words that followed pain stimulations after a several second delay, as had previously been described. This result is likely due to the unpredictable nature and variable positioning of non-pain items relative to painful ones. Future work that specifically controls for the ordering of pain-paired and non-pain experimental items could help to further clarify the temporal effects of memory impairment surrounding the brief experience of acute pain stimulation.

## Acknowledgments

This work was supported by the Department of Anesthesiology, University of Pittsburgh, School of Medicine. Further research funding and salary support was provided by a Mentored Research Training Grant (to KMV) from the Foundation for Anesthesia Education and Research (MRTG-CT 2-2017). Salary support for KMV during the different phases of this project came from the National Institutes of Health (T32GM075770 & K23GM132755). Additional financial support for KMV came from the NIH Clinical Loan Repayment program (L30GM120759). Subject recruitment was assisted by the University of Pittsburgh Clinical Translational Science Institute Research Participant Registry, a project supported by the National Institutes of Health (UL1 TR000005). The authors have no relevant financial or other conflicts of interest to disclose related to this work.

## Supplementary Materials

The supplementary materials are provided to allow the reader a more in-depth look at our data and analysis. Supplementary Figures 1 and 2 show RTs on a log_10_ scale, which is the transformed space in which the statistical analysis was performed. Supplementary Figures 3 and 4 show the RT distributions before and after transformation into log_10_ space, to display the distribution on which our parametric statistical analyses were performed. Supplementary Table 1 contains the hit rates and false alarm rates, for individual subjects’ Remember and Know responses, collected during the recognition testing portion of the experiment. This granularity allows verification of calculations used for our summary measures displayed in Figure 4, as well as the calculation of other metrics, if desired.

**Supplementary Figure 1.**
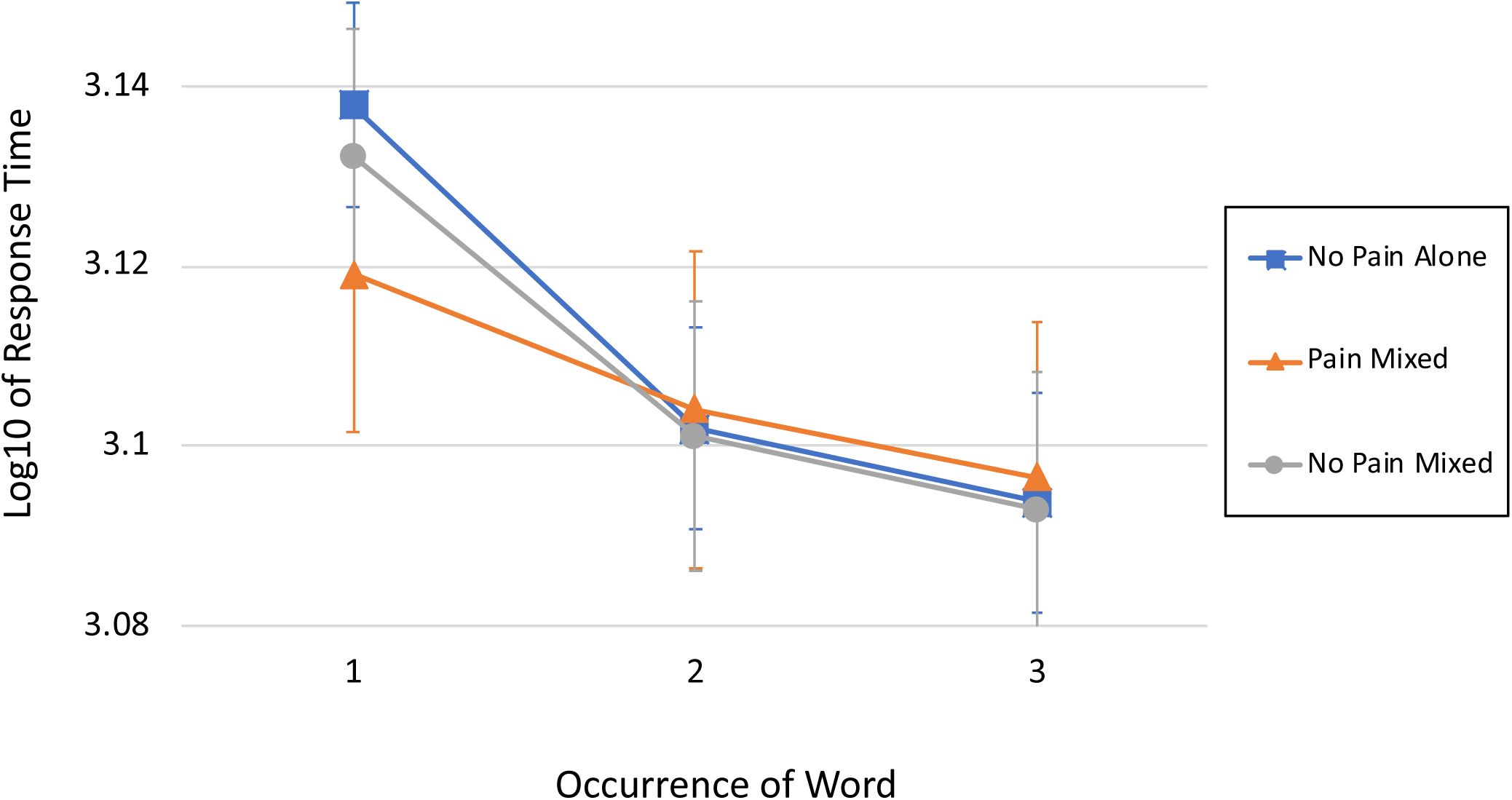
Common logarithm of response times over the three repetitions for each word-type during the Encoding portion of the experiment.

**Supplementary Figure 2.**
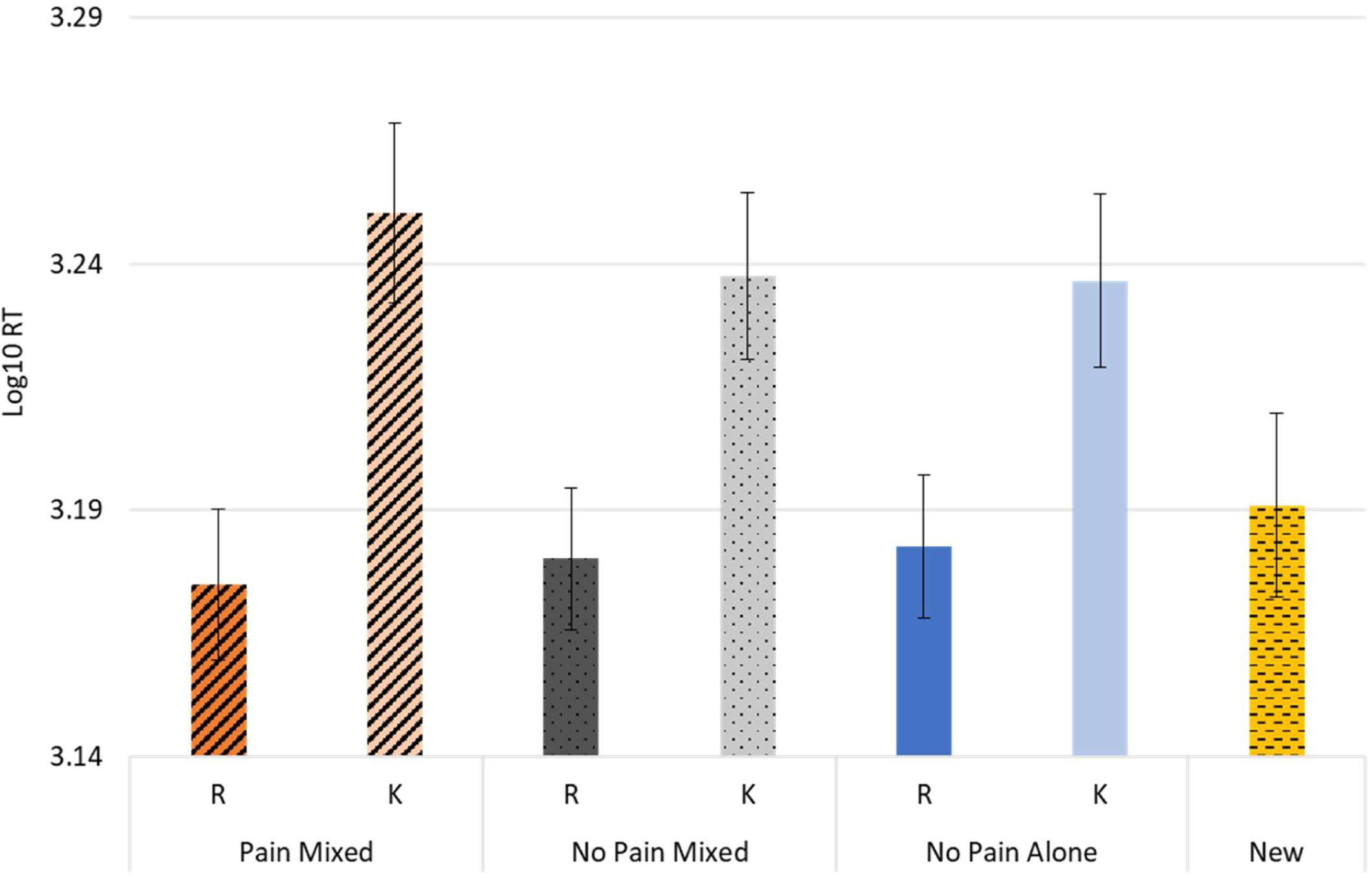
Common logarithm of response time data for Remember (R), Know (K), and New (N) responses during the RKN testing portion, including only correct responses. See text for discussion of significant differences.

**Supplementary Figure 3.**
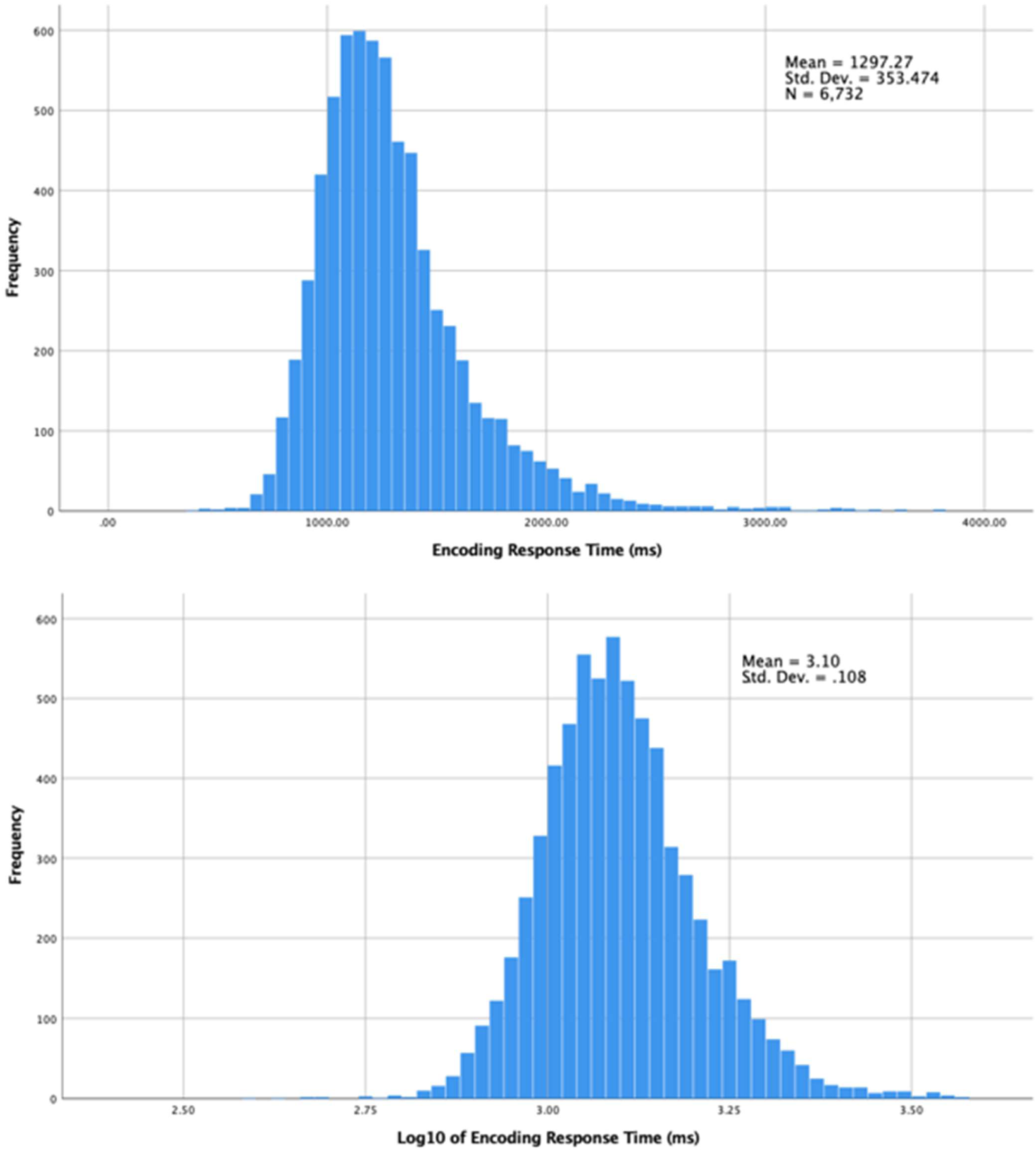
Histograms of encoding response times (top) and the same data transformed by the common logarithm (bottom).

**Supplementary Figure 4.**
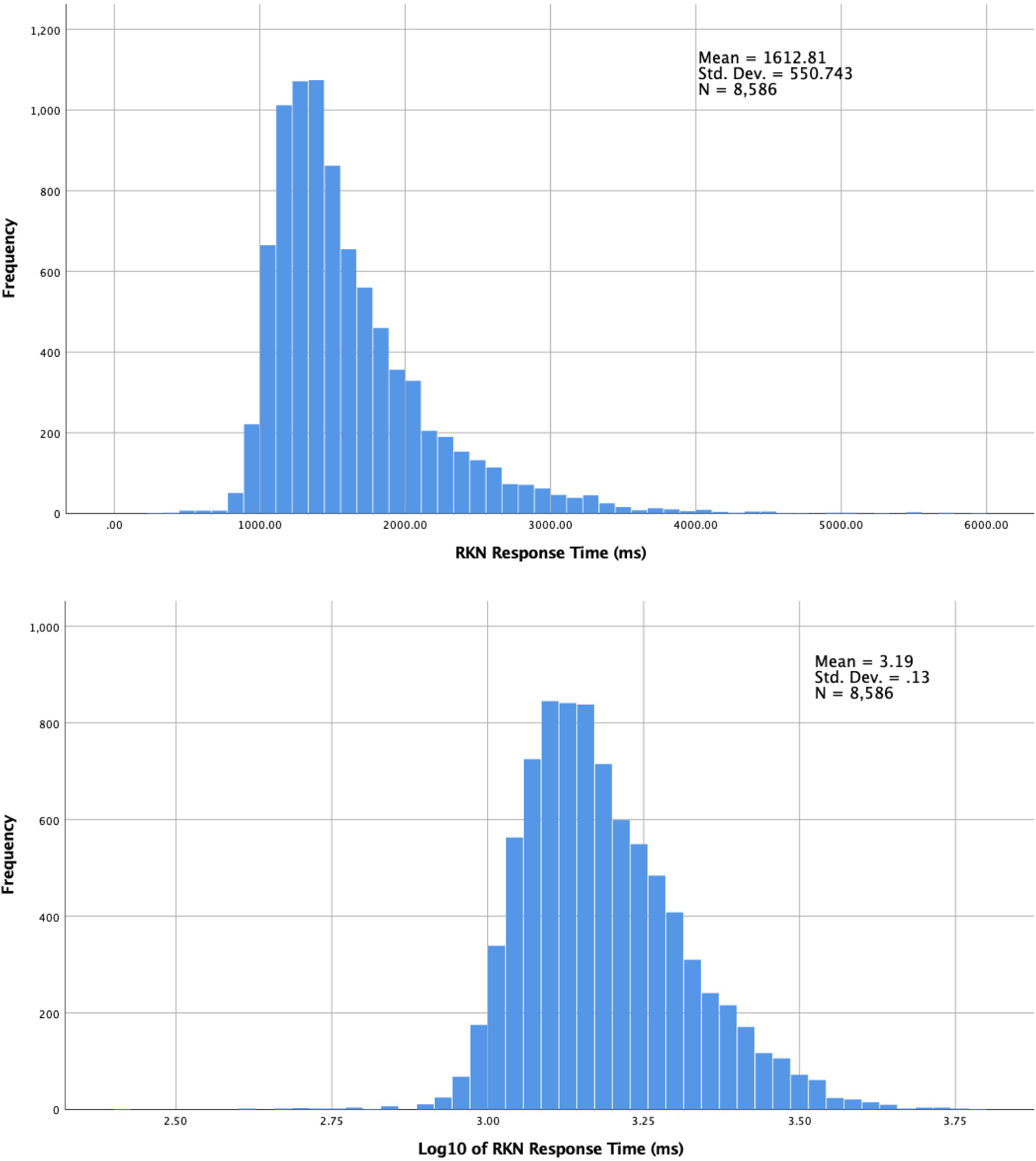
Histograms of RKN (Remember, Know, New) response times (top) and the same data transformed by the common logarithm (bottom).

**Supplementary Table 1.**
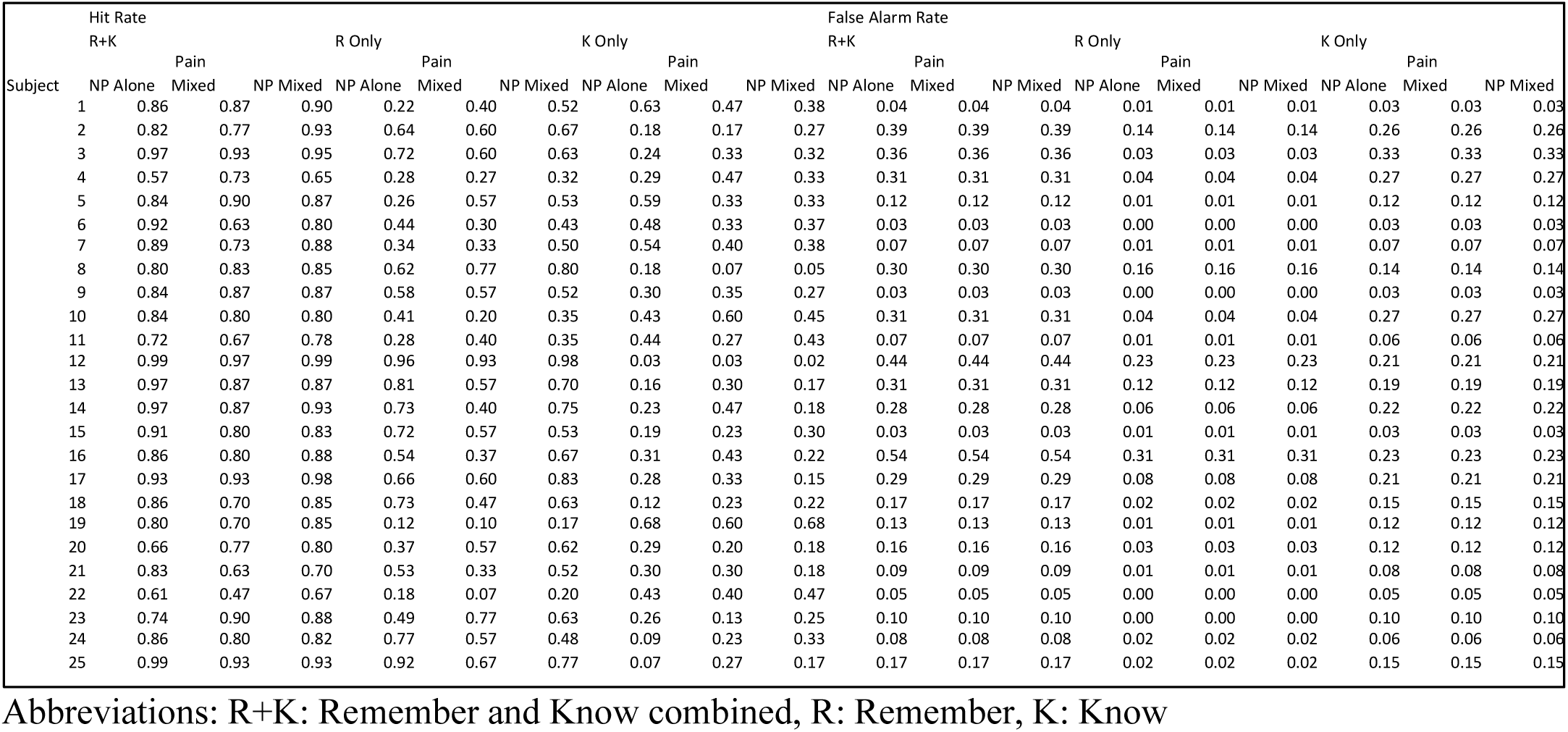
Hit Rates and False Alarm Rates for all word types and response types.

## References

Amado, S. (2000). Farkli dikkat düzeylerinin örtük ve açik bellek üzerindeki etkileri. [The effects of different levels of attention on implicit and explicit memory]. Türk Psikoloji Dergisi, 15(45), 41–56.

Baddeley, A. (2010). Working memory. Curr Biol, 20(4), R136–140. doi: 10.1016/j.cub.2009.12.014

Forkmann, K., Schmidt, K., Schultz, H., Sommer, T., & Bingel, U. (2016). Experimental pain impairs recognition memory irrespective of pain predictability. European Journal of Pain, 20(6), 977–988. doi: 10.1002/ejp.822

Kilb, A., & Naveh-Benjamin, M. (2015). The effects of divided attention on long-term memory and working memory in younger and older adults: Assessment of the reduced attentional resources hypothesis. In R. H. Logie & R. G. Morris (Eds.), Working memory and ageing (pp. 48-78, Chapter xv, 159 Pages): Psychology Press, New York, NY.

Liang, S., & You, X. (2003). The experimental dissociation between implicit and explicit memory tasks: The role of attention during encoding. Psychological Science (China), 26(4), 751–752.

Migo, E. M., Mayes, A. R., & Montaldi, D. (2012). Measuring recollection and familiarity: Improving the remember/know procedure. Conscious Cogn, 21(3), 1435–1455. doi: 10.1016/j.concog.2012.04.014

Mulligan, N. W. (1998). The role of attention during encoding in implicit and explicit memory. Journal of Experimental Psychology: Learning, Memory, and Cognition, 24(1), 27–47. doi: http://dx.doi.org/10.1037/0278-7393.24.1.27

Popov, V., & Reder, L. M. (2018). Frequency Effects on Memory: A Resource-Limited Theory. doi: https://osf.io/dsx6y/

Rajaram, S. (1993). Remembering and knowing: two means of access to the personal past. Mem Cognit, 21(1), 89–102. doi: 10.3758/bf03211168

Reder, L. M., Nhouyvanisvong, A., Schunn, C. D., Ayers, M. S., Angstadt, P., & Hiraki, K. (2000). A mechanistic account of the mirror effect for word frequency: a computational model of remember-know judgments in a continuous recognition paradigm. J Exp Psychol Learn Mem Cogn, 26(2), 294–320.

Rotello, C. M., & Zeng, M. (2008). Analysis of RT distributions in the remember-know paradigm. Psychonomic Bulletin & Review, 15(4), 825–832. doi: http://dx.doi.org/10.3758/PBR.15.4.825

Schwarze, U., Bingel, U., & Sommer, T. (2012). Event-related nociceptive arousal enhances memory consolidation for neutral scenes. J Neurosci, 32(4), 1481–1487. doi: 10.1523/JNEUROSCI.4497-11.2012

Vogt, K. M., Norton, C. M., Speer, L. E., Tremel, J. J., Ibinson, J. W., Reder, L. M., & Fiez, J. A. (2019). Memory for non-painful auditory items is influenced by whether they are experienced in a context involving painful electrical stimulation. Exp Brain Res. doi: 10.1007/s00221-019-05534-x

Wais, P. E., Mickes, L., & Wixted, J. T. (2008). Remember/know judgments probe degrees of recollection. J Cogn Neurosci, 20(3), 400–405. doi: 10.1162/jocn.2008.20041

Yonelinas, A. P., Aly, M., Wang, W. C., & Koen, J. D. (2010). Recollection and familiarity: examining controversial assumptions and new directions. Hippocampus, 20(11), 1178–1194. doi: 10.1002/hipo.20864

